# Simple controls exceed best deep learning algorithms and reveal foundation model effectiveness for predicting genetic perturbations

**DOI:** 10.1101/2025.01.06.631555

**Authors:** Daniel R. Wong, Abby S. Hill, Robert Moccia

## Abstract

Modeling genetic perturbations and their effect on the transcriptome is a key area of pharmaceutical research. Due to the complexity of the transcriptome, there has been much excitement and development in deep learning (DL) because of its ability to model complex relationships. In particular, the transformer-based foundation model paradigm emerged as the gold-standard of predicting post-perturbation responses. However, understanding these increasingly complex models and evaluating their practical utility is lacking, along with simple but appropriate benchmarks to compare predictive methods. Here, we present a simple baseline method that outperforms both state of the art (SOTA) in DL and other proposed simpler neural architectures, setting a necessary benchmark to evaluate in the field of post-perturbation prediction. We also elucidate the utility of foundation models for the task of post-perturbation prediction via generalizable fine-tuning experiments that can be translated to different applications of transformer-based foundation models to tasks of interest. Furthermore, we provide a corrected version of a popular dataset used for benchmarking perturbation prediction models. Our hope is that this work will properly contextualize further development of DL models in the perturbation space with necessary control procedures.

## Introduction

Understanding the phenotypic effects of genetic perturbations is a fundamental area of research in biology^1,2^. Due to the complex and interconnected circuitry of the human genome^3^, understanding perturbational effects and the repercussions this can have on other genes is a difficult task. Perturb-seq^4^ emerged as a tool to aid this discovery effort by measuring complex post-perturbation transcription profiles. This can be done at scale, facilitating the surge of data in the single-cell discipline^5–7^. Knowing what exact transcriptional changes result from different perturbations would be pivotal for both understanding human biology and progressing drug discovery. Since systematically perturbing genes can be an expensive and time-consuming endeavor, the idea of *in-silico*^8,9^ modeling and prediction becomes an attractive means of cost-efficient hypothesis generation. Hence, there has been a flurry of computational methods and especially DL ones to address this unmet need^1^.

Transformer-based models^10^ have captivated much of recent scientific efforts in a broad range of fields^11–14^. As excitement around large language model (LLM) success in Natural Language Processing grows^15^, so too do efforts of applying core concepts of LLMs to the “language” of biology^16–20^. Indeed the idea of “foundation models”^21^ is of particular excitement. Key to the idea of foundation models is the notion that pre-training on large amounts of data can yield model weights that capture foundational knowledge of the domain on which the model was trained. For single-cell biology, this can be a foundational understanding of complex gene networks learnable via self-supervision^22^. These foundation models can then be fine-tuned to adapt to related tasks, such as post-perturbation prediction. The promise is that foundational knowledge of fundamental biology can even bolster generalization to related tasks and new data via fine-tuning on much smaller datasets than the large corpus on which the foundation model was trained^21,23^. The prospect of potentially overcoming small data regimes via foundation models would indeed be a paradigm shift, especially in the field of computational biology and pharmaceutical research where data sets are often small, siloed^24,25^, and expensive to produce. Practically evaluating and understanding these exciting prospects in pharmaceutical use cases will be of great importance in advancing AI-driven drug discovery.

As a result of the excitement around foundation models in biology, there has been a large uptake of studies and methods that model single-cell biology via transformers^23,26–31^. With rising excitement and rapid development of new studies and methods, rigorous evaluation and proper attention needs to be given to understanding the performance and inner-workings of these black-box models, their limitations, and the biological context in which they can be most effective. There have been a growing number of studies in this area^32–40^, but much more analysis needs to be done to fully understand these models. Furthermore, as research in this area grows, we must standardize metrics and common-sense benchmarks to evaluate new methods^33,41–43^.

For the task of directly predicting the post-perturbation transcriptome, two DL methods achieved SOTA performance. The first is a graph-based learning method called GEARS^44^. The second is a transformer-based foundation model called scGPT^27^. In this study, we evaluate these two methods and compare them to a much simpler method that achieves SOTA performance, necessitating the need for more standard benchmarking. Through a series of control experiments that can be applied to any transformer-based foundation model, we also hope to introduce a general framework for evaluating both the utility of pre-trained foundation weights and the benefit of the transformer architecture for downstream tasks of interests, such as post-perturbation prediction performance. Through this study we hope to inform further model development for predicting single-cell perturbation responses.

## Results

To evaluate the utility of the leading DL methods for the post-perturbation prediction task, we analyzed their performance on three Perturb-seq datasets: Adamson^45^, Norman^46^, and Replogle K562 Essential^47^ (Methods). Due to the ease of data loading, many studies^27,28,33,43,44^ have recently used the ready-made datasets provided by the GEARS^44^ repository for training and evaluating models. For the popular Adamson dataset, which consists of three independent experiments, we noticed that the separate experiments were combined into one dataset. We wanted to keep consistent with the Norman and Replogle K562 Essential datasets, both of which had just a single independent experiment. Hence, we merged the GEARS annotations with the original metadata provided by the Adamson study authors, took the largest of the three constituent experiments, which focused on the unfolded protein response, and denoted this as the “Corrected Adamson UPR” dataset, which we use for our main analyses (Methods). In the process, we also discovered additional discrepancies between the original Adamson metadata and the annotations provided in the GEARS dataset, such as relabeling multiply perturbed cells in the “epistasis” experiment as controls. While not the focus of the present study, we provide a revised version of the GEARS Adamson dataset (consisting of all three experiments and realigned with the original author guide calls) as a convenience to the community (Table 1).

**Table 1:**
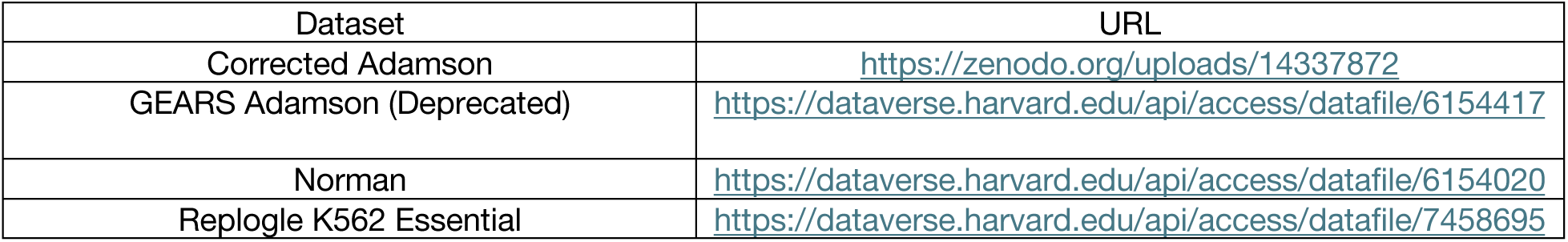
dataset locations.

| Dataset | URL |
| --- | --- |
| Corrected Adamson | https://zenodo.org/uploads/14337872 |
| GEARS Adamson (Deprecated) | https://dataverse.harvard.edu/api/access/datafile/6154417 |
| Norman | https://dataverse.harvard.edu/api/access/datafile/6154020 |
| Replogle K562 Essential | https://dataverse.harvard.edu/api/access/datafile/7458695 |

We fine-tuned a pre-trained scGPT Human model^27^ using the default parameters that the authors provided for fine-tuning on Perturb-seq data. We also trained GEARS using default parameters. For evaluating prediction of the transcriptome for held-out perturbations, Roohani et al.^44^ proposed measuring the Pearson correlation between the actual and predicted expression. As a further metric, Pearson correlation can be computed over just the top 20 differentially expressed (DE) genes for a perturbation (denoted Pearson DE).

Cui et al.^27^ further adapted the Pearson and Pearson DE scores by subtracting control expression to get a more granular metric that measures similarity between differences called Pearson delta (PD). Similarly, this can be computed for just the top 20 differentially expressed genes and is denoted as Pearson DE Delta (PDED) (Methods). Supplemental Figure 1 provides a pictorial illustration of the different standard metrics. Since delta metrics provide a more informative readout of transcriptional changes^27^, we focus our analyses on these scores.

### CRISPR-informed mean is a SOTA predictor

As a simple competitive method for predicting the effects of held-out perturbations on transcriptomic expression, we calculated the mean expression for each gene over the perturbed cells in the training set and stipulated this as a prediction for the test set. Other recent studies have also proposed simple baseline mean variants as competitive methods to more complex regression and even DL-based methods^34–40^. It is important to note that the training set perturbations were completely disjoint from the test set perturbations. We denote this model as the “mean model” (Methods). For the PD metric, the mean model outperformed GEARS for all three datasets (difference of means = 0.07, p-value = 0.01) and outperformed scGPT for all datasets (difference of means = 0.10, p-value = 1.6×10^-4^) (Figure 1). For the PDED metric, the mean model exceeded GEARS for 2/3 datasets (difference of means = 0.03, p-value = 0.38 across all datasets), while scGPT outperformed the mean model for 2/3 datasets (difference of means = 0.04, p-value = 0.14) (Figure 1). Statistical significance was reached for differentiating the mean model from both GEARS and scGPT for the PD metric but not for the PDED metric. Similarly, mean as a superior predictor to GEARS and competitive with a different LLM is reported by Märtens et al.^36^ for the Replogle dataset.

**Figure 1:**
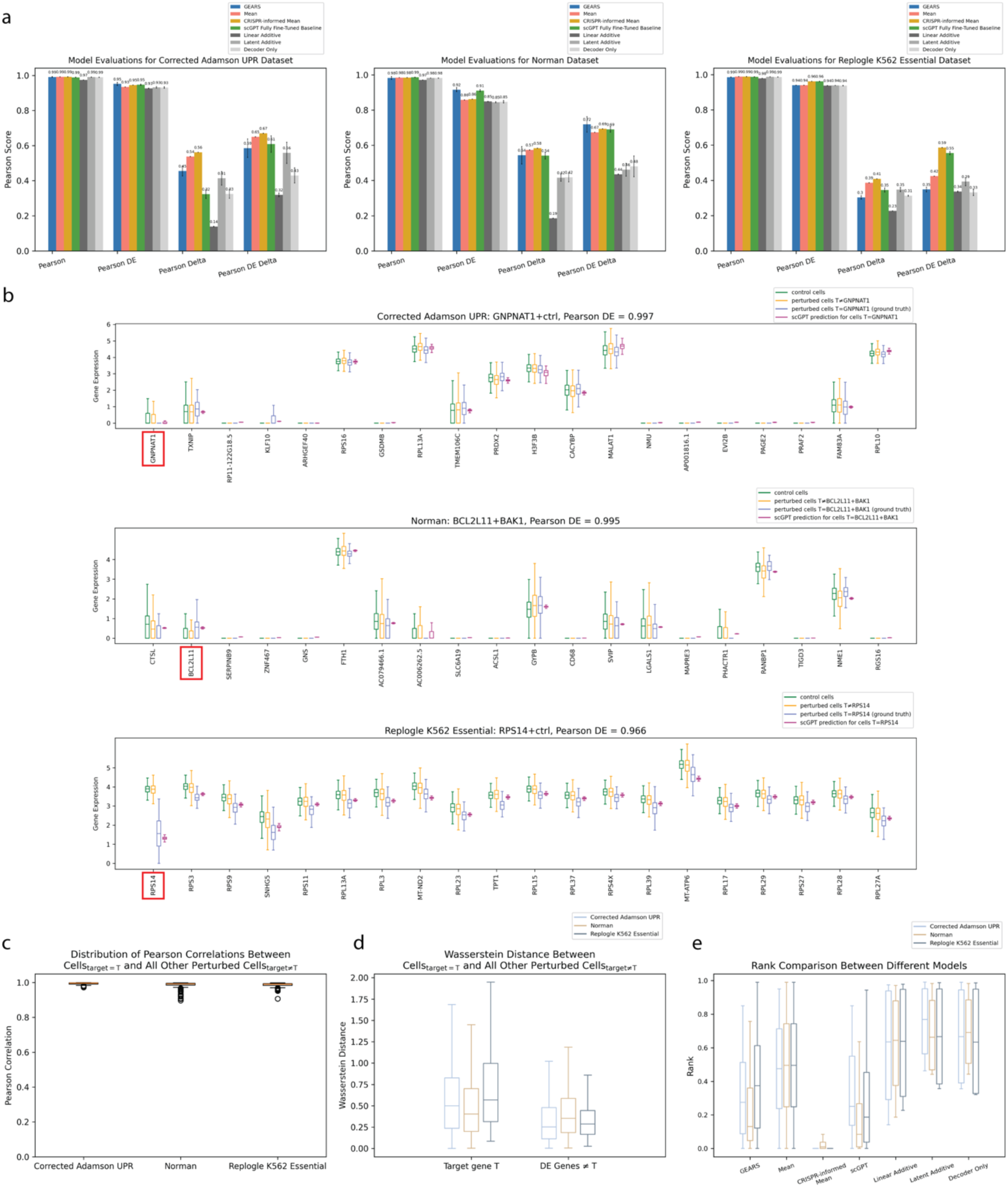
**CRISPR-informed mean as SOTA predictor.** (a) Performance bar charts on held-out test set for different models. Error bars span plus and minus one standard deviation of 10 independently trained models applied to the same test set. (b) Boxplots of top 20 DE genes for a subset of scGPT’s high performing held-out perturbation conditions for each dataset. Target gene T is boxed in red when T is among the top 20 DE genes. The title of each plot denotes the dataset, perturbation condition, and scGPT’s Pearson DE score for predicting the expression of cells_target=T_. In the figure legend “T≠condition” denotes all perturbed cells_target≠T_ in the dataset. “T=condition” denotes all cells_target=T_. (c) Boxplots of Pearson correlations between average expression of cells_target=T_ and average expression of perturbed cells_target≠T_ repeated over each target T in the whole dataset. Pearson correlation was calculated over all genes for all perturbations in the dataset. (d) Boxplots of Wasserstein distances. Left: boxplots of Wasserstein distances between (1) expression of T in cells_target=T_ and (2) expression of T in perturbed cells_target≠T_. Right: boxplots of Wasserstein distances ∀ gene D≠T ∈ DE genes between (1) expression of D in cells_target=T_ and (2) expression of D in perturbed cells_target≠T_. Boxplots are computed over all perturbations in the dataset. The differences between the left and right distributions were all statistically significant with the following p-values for a two-sided t-test: Corrected Adamson UPR = 3.6×10^-13^, Norman = 4.5×10^-5^, Replogle K562 Essential = 4.3×10^-173^. (e) Rank comparisons between different methods over the held-out perturbation test set. A rank of 0.0 indicates a perfect score, and a rank of 0.5 is the expected score from a random prediction. For held-out perturbation with target=T, the reference gene expression used to compute rank was the average of the ground truth expression of cells_target=T_ over all genes. Since the prediction of the mean model is the same for all perturbations, the cosine similarity to reference is the same for all perturbations and results in a random rank. For each model type, we used the best-performing model from the ten independently trained models that achieved the highest PDED performance on the test set.

We visualized scGPT’s top 20 highest-scoring perturbations for differentially expressed genes (Figure 1B). From this visual representation, we observed why the mean expression of perturbed cells is a competitive predictor: a single perturbation with target=T does not induce much change in gene expression of cells with target = T (denoted cells_target=T_) relative to the expression of all other perturbed cells with target ≠ T (denoted cells_target≠T_) (Figure 1C). Hence, simply predicting the average of perturbed cells_target≠T_ results in a profile similar to the average over cells_target=T_ (the ground truth). For the highly performant perturbations in Figure 1B, the distribution of T’s expression in cells_target=T_ is dissimilar to the distribution of T’s expression in all other perturbed cells_target≠T_. In contrast, cells_target=T_ and all other perturbed cells_target≠T_ have similar distributions for DE genes ≠ T. Quantifying this observation with the Wasserstein distance across all perturbations, we found that this trend held for both CRISPRi datasets (Adamson and Replogle K562 Essential) and (to a lesser extent) for the CRISPRa Norman dataset (Figure 1D).

Since predicting expression of the target gene is trivial for an effective CRISPRi experiment, we constructed a stronger predictor that exploits this biological knowledge. As a more informed but simple method, we implemented a model we denote as the “CRISPR-informed mean model” that leverages prior information about Perturb-seq and how CRISPRi should reduce transcriptomic expression to near 0 and how CRISPRa should enhance expression. This model predicts the mean of all perturbed cells in the training set for all genes except for the target gene T. For CRISPRi the model predicts 0 expression for T. For CRISPRa the model doubles the training set’s mean expression of T and stipulates this as its prediction for T (Methods). Since CRISPRi in practice hardly eliminates expression in full^48^ and CRISPRa has a wide range of enhancement based on the biological context^49^, this model is an overly simplistic approximation. However, this simple model that leverages known biological priors resulted in statistically significant SOTA performance over front-running DL models (Figure 1A). For the PD metric, the CRISPR-informed model outperformed both GEARS (difference of means = 0.08, p-value = 9.3×10^-4^) and scGPT (difference of means = 0.11, p-value = 8.1×10^-6^) for all three datasets. For the PDED metric, the CRISPR-informed model outperformed GEARS on 2/3 datasets (difference of means = 0.10, p-value = 0.002) and outperformed or was equivalent to scGPT (difference of means = 0.03, p-value = 0.03) on all datasets. For the Norman dataset, which used CRISPRa, the CRISPR-informed model lagged slightly behind GEARS (PDED difference = 0.03) and was near equivalent with scGPT (PDED difference = 0.002).

To help standardize the field of perturbational modeling, recent work from Wu et al.^43^ proposed other simple neural baseline models by which to measure new ML methods: Linear Additive, Latent Additive, and Decoder Only. These models utilized simple feed-forward neural architectures. We compared CRISPR-informed mean to these other models and found that CRISPR-informed mean was more performant than all other simple neural models for all datasets (Figure 1A). We also assessed performance of all models using the rank metric, a new metric proposed by Wu et al.^43^ for assessing post-perturbation prediction that focuses on triaging perturbations (Figure 1E). We found similar results, with CRISPR-informed mean surpassing all DL models for all datasets. Rank improvement ranged from 4.7-213.9x better than GEARS and 3.9-155.4x better than scGPT depending on the dataset. CRISPR-informed mean also had substantially better rank than all other simple neural baseline models for all datasets.

We also implemented four-fold cross-validation for all datasets and found similar results indicating the competitiveness of the CRISPR-informed model (Supplemental Figure 2). Even though this model was better on average than all DL models, there were perturbation conditions in which the DL models were superior to the simple CRISPR-informed model, but for most conditions the CRISPR-informed model was the higher performer for all datasets (Supplemental Figure 3). Furthermore, in a more granular analysis of which genes were the worst and best performing, we observed variable performance and a large overlap in top performing genes between the CRISPR-informed model and scGPT for all three datasets (Supplemental Figure 4).

### Neither foundation weights nor transformer attention provide a competitive advantage

Along with establishing proper benchmarks, it is important to understand the feasibility of foundation models transferring learned knowledge derived from pre-training into improving related tasks^21^. Even more important than this is a general framework that can test this feasibility for the sake of assessing future model development. To determine whether the single-cell foundation model weights could aid with the related post-perturbation prediction task, we compared a fully fine-tuned scGPT model, which had all foundation weights loaded from the Human CP model, to different fine-tuned scGPT models that selectively withheld different foundation weights prior to fine-tuning (Figure 2A). For the first weight-loading permutation, we trained a model that had an identical architecture to the fully fine-tuned scGPT model but initialized with random weights instead of leveraging any pre-trained weights (scGPT no-pretraining). We found that the metrics on the held-out test set were near identical across datasets, with no statistically significant difference in performance between the fully fine-tuned foundation model and the model trained from random weight initialization. The difference in PD means across all three datasets between the fully fine-tuned model and scGPT no-pretraining (ten independently trained models per dataset) was only 0.004 (p-value = 0.89) (Figure 2B). For PDED, this difference of means was only 0.01 (p-value = 0.75). For both metrics, we could not reject the null hypothesis that the means are identical, and we attributed the differences in mean largely to chance.

**Figure 2:**
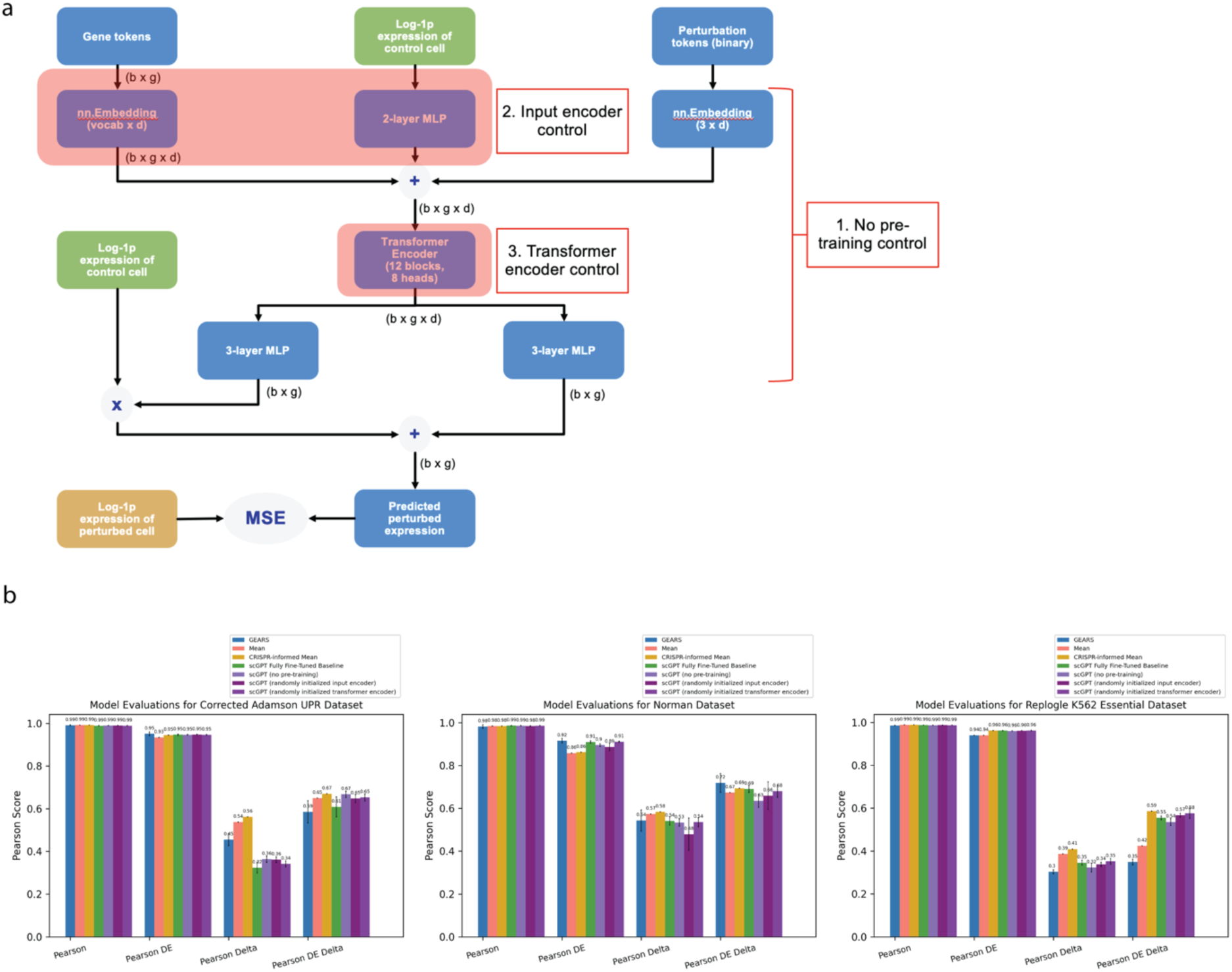
**Performance of weight and architecture permutations.** (a) scGPT model architecture for the Perturb-seq task. Here we denote 3 different ways to selectively use the foundation weights. (1) No-pretraining, i.e. not using them at all and initializing all weights randomly (2) loading the foundation input encoder weights and initializing all other weights randomly and (3) loading the foundation transformer encoder weights and initializing all other weights randomly. (b) Performance bar charts on held-out test set for different baseline models. Error bars span plus and minus one standard deviation of 10 independently trained models.

For the second weight-loading permutation, we loaded all the foundation weights minus the learned input encoder weights (withholding the learned gene embedding values and the expression encoder). We observed a PD difference of means = 0.01 (p-value = 0.66), and a PDED difference of means = 0.006 (p-value = 0.69). For the last permutation, we loaded all the foundation weights minus the transformer block weights. The self-attention mechanism, which is the key innovation that defines transformers^10^, is completely contained within this transformer block. A similar trend held, with a PD difference of means = 0.006 (p-value = 0.80), and a PDED difference of means = 0.02 (p-value = 0.22). scGPT with a randomly initialized transformer did slightly better on average than the fully fine-tuned scGPT with all foundation weights loaded prior to fine-tuning (but did not achieve statistically significant improvement). As an additional experiment, we ensured the persistence of foundation weights by fine-tuning the pre-trained scGPT foundation model using a popular alternative to full fine-tuning for LLMs called Low Rank Adaptation (LoRa)^50^, which relies on freezing the foundation weights and learning rank decomposition matrices. We found that the LoRa models slightly underperformed against the fully fine-tuned models (Supplemental Figure 5), which seems consistent with LoRa applied in other contexts^51,52^.

To answer the question of whether the transformer architecture provides a competitive advantage over simpler architectures, we trained a competing model that is identical to scGPT, but without the transformer encoder (denoted as “Simple Affine”, Figure 3A). After training the Simple Affine model using random weight initialization, we found that this model was competitive with the scGPT model without pre-training with a PD difference of means = 0.02 (p-value = 0.49) and PDED difference of means = 0.02 (p-value = 0.24) indicating that the transformer architecture itself added little utility over a simpler architecture for post-perturbation prediction (Figure 3B). Despite having near identical performance results, scGPT trained 19-20x slower than Simple Affine depending on the dataset (Figure 3C) and was about 1.6x the size of Simple Affine (Figure 3D). Instead of simply removing the transformer block, we also performed an experiment where we replaced the transformer block with a simple multilayer perceptron (MLP) with the same number of layers and observed similar results (Supplemental Figure 6).

**Figure 3:**
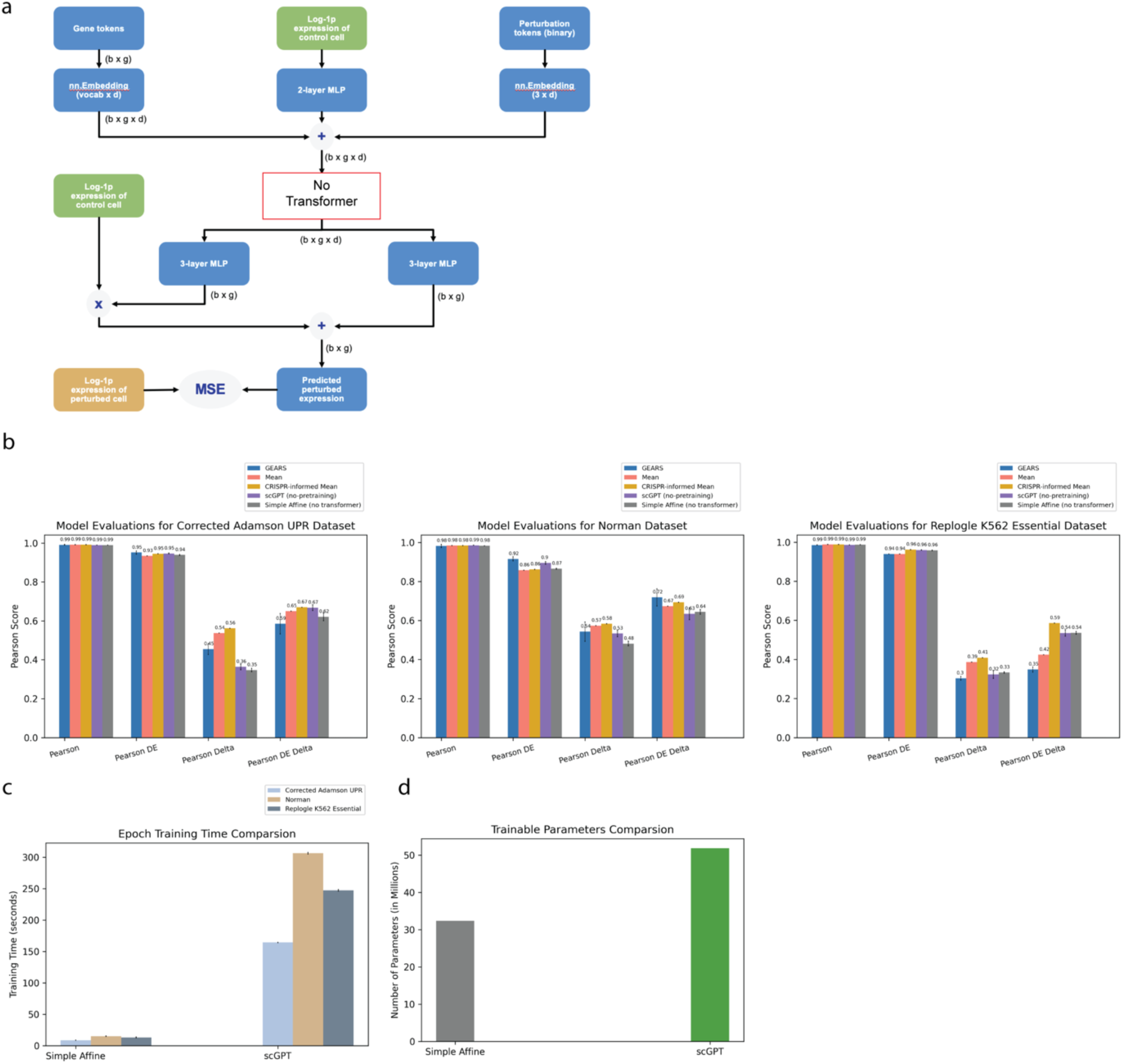
**Simple Affine model is competitive with transformer-based model.** (a) Architecture of the Simple Affine model. (b) Performance bar charts on held-out test set for different baseline models. Error bars span plus and minus one standard deviation of 10 independently trained models. (c) Bar charts comparing Simple Affine and scGPT average training time per epoch for each dataset on one Nvidia H100 GPU. (d) Bar charts comparing Simple Affine and scGPT model sizes.

For the sake of consistency with other studies, we also report the corresponding results when we trained models on the GEARS version of the Adamson dataset (Supplemental Figure 7). We also report results when we trained models on the entirety of the corrected Adamson dataset, utilizing all three of its constituent experiments (Supplemental Figure 8).

## Discussion

Predicting the post-perturbation transcriptome is a difficult task that has many opportunities for further development. At least for this task, current DL and foundation models do not yet demonstrate meaningful improvement over simple but inadequate methods. As new computational models are being developed, there must be an adoption of standard benchmarks and methods like CRISPR-informed mean to properly assess new methods. Although a SOTA predictor, a CRISPR-informed mean is not sufficient for most drug discovery use-cases because it does not contextualize gene to gene interactions, cannot be used in zero-shot settings because different datasets often have different means, and cannot possibly model nuanced changes in expression. This sort of baseline model cannot be applied to experiments where the target is unknown, which is common in drug discovery settings especially with small molecule perturbations. Since Perturb-seq is a destructive process and cannot result in true pairings between pre and post perturbation cells^4^, predicting the post-perturbation transcriptome is innately a prediction of a statistical distribution. This fact, coupled with the fact that CRISPR perturbations result in subtle and minimal transcriptional changes across the entire transcriptome (as evident by high Pearson correlations between cells_target=T_ and cells_target≠T_, Figure 1C), reinforces how statistical baselines like mean (and related but more biologically-sensible versions of mean) are appropriate metrics that need to be included in all post-perturbation prediction benchmarks. Despite being similar, the CRISPR-informed mean model often substantially outperformed the standard mean model across all datasets for rank and Pearson metrics (Figure 1A, 1E), necessitating its inclusion in all future benchmarking. This also reveals the sensitivity of current metrics to changes in prediction of just one or two genes, which is essential for a data type in which expression is not that differentiable between perturbations. Models that are to exceed mean or variants of it must also be sensitive enough to detect and learn from subtle transcriptional changes, while also being general enough to reflect universal gene relationships. For the fundamental question of how perturbing one gene affects other genes, any prospective model that underperforms against this simple model, which has no possibility of reflecting gene to gene interactions, cannot be regarded with high confidence. Hence the CRISPR-informed mean can serve as a benchmark to inform whether any new models developed can be of practical use in discerning subtle changes in biology.

We appreciate the many community efforts towards increased standardization and benchmarking, such as the simple neural models presented by Wu et al^43^. The underlying themes of such work are commendable for improving the field of perturbation modeling via standardized benchmarks and reproducible science. Since CRISPR-informed mean is both more performant than these neural baseline models (Figure 1A) and easier to implement, we advocate for the adoption of CRISPR-informed mean as the simplest and most performant benchmark for post-perturbation prediction tasks over all DL and non-DL baselines we studied. As new proposed ML methods get increasingly more complex, we must ensure that such designs merit the extra complexity, and right-size their performance with interpretable and sensible baselines. We expect that leveraging known biology directly in model development beyond the simplistic design of CRISPR-informed mean will be an avenue of further research. As important as having standardized model benchmarks, standardized datasets are just as essential for furthering community development of perturbation models. By providing a corrected version of a popular dataset used by the ML community, we hope to avoid any future analyses based off erroneous data labeling, encourage greater scrutiny for pre-processing data and checking the work of others, and facilitate easy adoption of a useful dataset ready for ML analyses.

The idea of attention is perhaps a promising solution to complex gene to gene relationships. However, as with most transformer-based models, copious and diverse training data is required, for which Perturb-seq may not currently have enough to model complex post-perturbation responses^6^. Hence, as important as standardized benchmarking and competing models are, so too is the need for the generation of diverse, large perturbational datasets if we are to leverage the full utility of transformer models and replicate their high performance in the NLP space fueled by large datasets^53^. Furthermore, as more investigation continues in developing transformer-based foundation models for application in the single-cell space, proper attention controls like the ones demonstrated in Figure 2 are also needed to assess whether these foundation models can truly translate foundational biological knowledge to pertinent tasks of interest. For this specific task of post-perturbation prediction, if the foundation model had proper gene-to-gene attention, then it was not advantageous over simpler models without self-attention. If the proper attention was not present to begin with, then it certainly was not learned during fine-tuning (such that it could substantially benefit performance compared to simpler models), nor would we expect it to be robustly learned given the small dataset sizes. The control experiments we studied here can be applied in general to various use cases of fine-tuning large foundation models. These controls will be invaluable in assessing if such models can provide foundational knowledge for performance improvement of downstream tasks. Perhaps these should be standard practices and sanity checks whenever one proposes a new foundation model advertising increased utility to related tasks. Both procedures of withholding foundation weights prior to fine-tuning and stripping architectures of their attention mechanisms should drastically reduce performance if a model is indeed properly translating learned foundational knowledge. Even though the transformer-based foundation models underperformed simpler models, it is worth noting that transformer-based models may yield both better gene correlations across perturbations and better PD scores for perturbations across genes compared to non-attention based (but otherwise equivalent) model architectures. This advantage may be slight with magnitude varying by dataset and by whether pre-training was used (Supplemental Figure 9). This provides a more optimistic but tempered outlook on the promise of transformers and large pretraining applied to single cell perturbation modeling.

Since larger models tend to require more data to perform well and avoid overfitting, it is essential to check if simpler architectures can deliver similar performance while being more efficient (Figure 3), especially in a data-sparse field like Perturb-seq. If transformers are employed, we must ensure that whatever learned self-attention merits the extra complexity and compute costs. Since data is expensive to generate, another opportunity for improved performance is to leverage known biological knowledge directly into modeling. As observed with the large performance gains of CRISPR-informed mean over simple mean, introducing biological priors can be a small but beneficial modification. Foundation models also present a similar opportunity of injecting known biology, but we need not forgo other simpler ways to leverage biological knowledge in computational models. We imagine that a combination of both would be the most advantageous, necessitating close interdisciplinary collaboration between computer scientists and biologists to achieve fundamental understanding of foundation models and ways they can be adapted to the biological domain.

In addition to further exploring foundation models, more attention needs to be paid towards metric development. Current metrics that evaluate predicted gene expression, like mean-squared-error and Pearson correlation and its variants, all do not address how useful model predictions are to broader pharmaceutical goals. Knowing the exact expression of different genes in response to perturbation is undoubtedly useful as an initial readout for early drug discovery, but greater downstream questions remain. Can these predictions inform experimental design and streamline costs by proposing which perturbations to test in which model systems? Are these predictions valid across different cell lines or organisms? Can they determine which perturbations directly affect disease biology and have therapeutic promise? These are all larger questions that current metrics are insufficient to address. We hope that future work in this field will investigate biologically useful metrics as well as model design.

In conclusion, we present three examples of simple controls: CRISPR-informed mean, selective withholding of foundation weights prior to fine-tuning, and removal of self-attention. Collectively, these controls set new gold standards for performance, shed light on the current state of the field, and provide proper checks for the further development of computational models applied to single-cell biology. We also hope that our work revising a popular dataset would serve as a case study for promoting heightened attention towards data preprocessing for ML training. Although we are optimistic about the potential for DL and foundation models to transform perturbational analyses in single-cell biology, we advocate for the adoption of similar controls and a deeper understanding of increasingly complex models.

## Methods

### Statistical Tests

For all statistical tests comparing two distributions, we used a two-sided t-test as implemented by SciPy’s stats.ttest_ind (version 1.10). The null hypothesis was that the means of the two distributions are equal. We used a significance cutoff of p-value < 0.05 to reject the null hypothesis. For comparing different model performances (model A and model B) across datasets, the sample size was 30 for each model result (three datasets with ten independently trained models per dataset). Each independently trained model was applied to the same fixed test set. The difference of means reported in the Results section was simply the mean of the performance results from model A (n=30) – the mean of the samples from model B (n=30).

### Dataset Curation

We downloaded the GEARS Adamson, Norman, and Replogle K562 Essential datasets from the Harvard Dataverse using the GEARS interface in June 2024. We did not perform any additional preprocessing steps on the provided data, which had log-transformed expression matrices. The Adamson and Replogle dataset were single-target CRISPRi, while the Norman dataset was multi-target CRISPRa. All datasets consisted of K562 cells. For the Replogle K562 Essential dataset, we filtered out cells_target=T_ for all T that was not present in the transcriptomic readout.

We also provide the Corrected Adamson dataset. Briefly, the original pre-processing in the GEARS repository contained perturbed cells mislabeled as control cells. We downloaded the original source data from the NCBI Gene Expression Omnibus (GSE90546) to generate AnnData files for each of the three experiments included in the submission. The cell metadata was then compared back to what was documented in the curation of the same dataset provided by the GEARS authors. This revealed a substantial number of cells where the condition label disagreed. Many of these mismatches were the result of cells mislabeled as “ctrl” (i.e. control) in GEARS, but with gene perturbations documented in the cell_identities.csv files provided in the original Adamson et al. paper^45^. We corrected the labels of these cells mislabeled as control and created a new AnnData object for release. See: https://github.com/pfizer-opensource/perturb_seq/tree/main/dataset_correction/ for all source code used to generate the corrected dataset. All datasets and URLs are provided in Table 1.

### Dataset Splitting

For all datasets, we constructed the training, validation, and test sets such that any unique perturbation was randomly assigned to only one of the sets. Dataset sizes and perturbation counts are shown in Supplemental Figure 10. Some perturbations in the Norman dataset utilized combinations of multiple genes (e.g. a single perturbation could have multiple gene targets A and B for A≠B). Combination perturbations are counted as unique perturbations and are each only present in one of the three sets. The perturbation splits for each dataset can be found at https://github.com/pfizer-opensource/perturb_seq/tree/main/splits/.

### DE Gene Selection

Selection of DE genes was identical to the methods used in Cui et al.^27^ and Roohani et al.^44^ For a perturbation P, we used Scanpy’s rank_genes_groups (version 1.9.3) function grouped by the perturbation condition. We used all control cells in the dataset as the reference for the t-test. We performed DE gene selection independently for each perturbation.

### Mean models

The mean model computed the mean expression of every gene across all the perturbed cells in the training set. The model returned the mean vector m as its prediction for all cells regardless of the query perturbation.

E_ij_ = expression of cell i for gene j

C_p_ = set of perturbed cells in the training set

m_j_ = value for gene j in mean vector

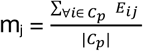

For the CRISPR-informed mean model, the model returned m as its prediction as defined by the following:

t = target gene

if CRISPRi:

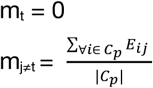

if CRISPRa:

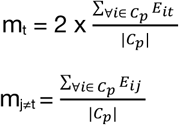

### DL Models

We trained The GEARS model (version = 0.1.2) using the default hyperparameters presented in the original study, a batch size of 64 cells, and 20 epochs of training. We trained scGPT (version = 0.2.1) fully fine-tuned by loading the published Human CP foundation model weights and fine-tuning with the same hyperparameters and setup as the original study’s fine-tuning on Perturb-seq data: batch size of 64, 15 epochs, learning rate of 0.0001, embedding size of 512, 12 transformer blocks, and 8 attention heads. For the scGPT model trained with random weight initialization (Figure 2), we did not load the pre-trained Human CP foundation weights and instead randomly initialized all model weights using PyTorch’s default weight instantiation for all weights in the model. We applied the same training procedure and hyperparameters as scGPT fully fine-tuned. For the various models that selectively loaded subsets of the pre-trained foundation weights from Human CP and initialized the rest at random (Figure 2), we also applied the same training procedure as scGPT fully fine-tuned. For all DL models, we trained 10 independent models from 10 different random initializations and kept the train-validation-test split constant across all models. We selected the final model weight for a given training run according to the default scheme for each method as follows. For scGPT, we selected the model weights that resulted in the highest Pearson correlation over all genes in the validation set. For GEARS, we selected the model weights that resulted in the lowest mean squared error over DE genes. We trained all models using a single Nvidia H100 GPU.

### Training other simple neural baseline models

For the simple baseline neural models presented by Wu et al.^43^ (Linear Additive, Latent Additive, and Decoder Only) we trained these models using the default hyperparameters, a batch size of 64, and 15 epochs. We used the same training, validation, and test sets as the DL models. During a training run, we selected the model weights according to the default validation selection by choosing the ones that had the lowest mean squared error loss between predicted and actual expression over the validation set.

### Performance Metrics

Roohani et al.^44^ and Cui et al.^27^ collectively proposed four Pearson metrics for assessing how well the post-perturbation transcriptome can be predicted. They can be described as follows:

t = target gene

C_c_ = set of control cells

C_t_ = set of perturbed cells_target=t_

G = set of all genes in the transcriptome

G_DE_c_ = set of 20 top differentially expressed genes for C_c_

G_DE_t_ = set of 20 top differentially expressed genes for C_t_

E_ij_ = actual expression of cell i for gene j

Ê_ij_ = predicted expression of cell i for gene j

C_mean_ = mean of control cells (all genes) = 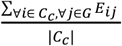

C_mean__DE = mean of control cells (just DE genes) = 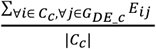

y_t_ = average of true expressions for C_t_ (all genes) = 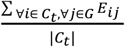

ŷ_t_ = average of predictions for C_t_ (all genes) = 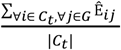

y_t_DE_ = average of true expressions for C_t_ (just DE genes) = 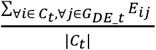

ŷ_t_DE_ = average of predictions for C_t_ (just DE genes) = 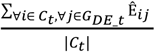

Pearson(t) = Pearson correlation(ŷ_t_, y_t_)

Pearson DE(t) = Pearson correlation(ŷ_t_DE_, y_t_DE_)

Pearson Delta(t) = Pearson correlation(ŷ_t_ - C_mean_, y_t_ - C_mean_)

Pearson DE Delta(t) = Pearson correlation(ŷ_t_DE_ - C_mean_DE_, y_t_DE_ - C_mean_DE_)

For summary Pearson metrics reported in the Results section, we averaged the Pearson scores over every target:

P = the set of all perturbations

Summary Pearson Metric = 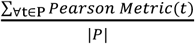

For the rank metric^43^, the rank score of a perturbation with target=t is calculated as the index of the average prediction for t in a sorted list of all average predictions for all perturbations (including target=t). The list is sorted from highest cosine similarity (to the actual average expression of cells with target=t) at the lowest index to lowest cosine similarity at the highest index. Lower ranks indicate better prediction, with a perfect prediction having rank = 0.0 and expectation from random having rank = 0.5.

## Data and Code Availability

All datasets are public and can be found at the links provided in Table 1.

All source code is available at: https://github.com/pfizer-opensource/perturb_seq

## Supplemental Figures

**Supplemental Figure 1:**
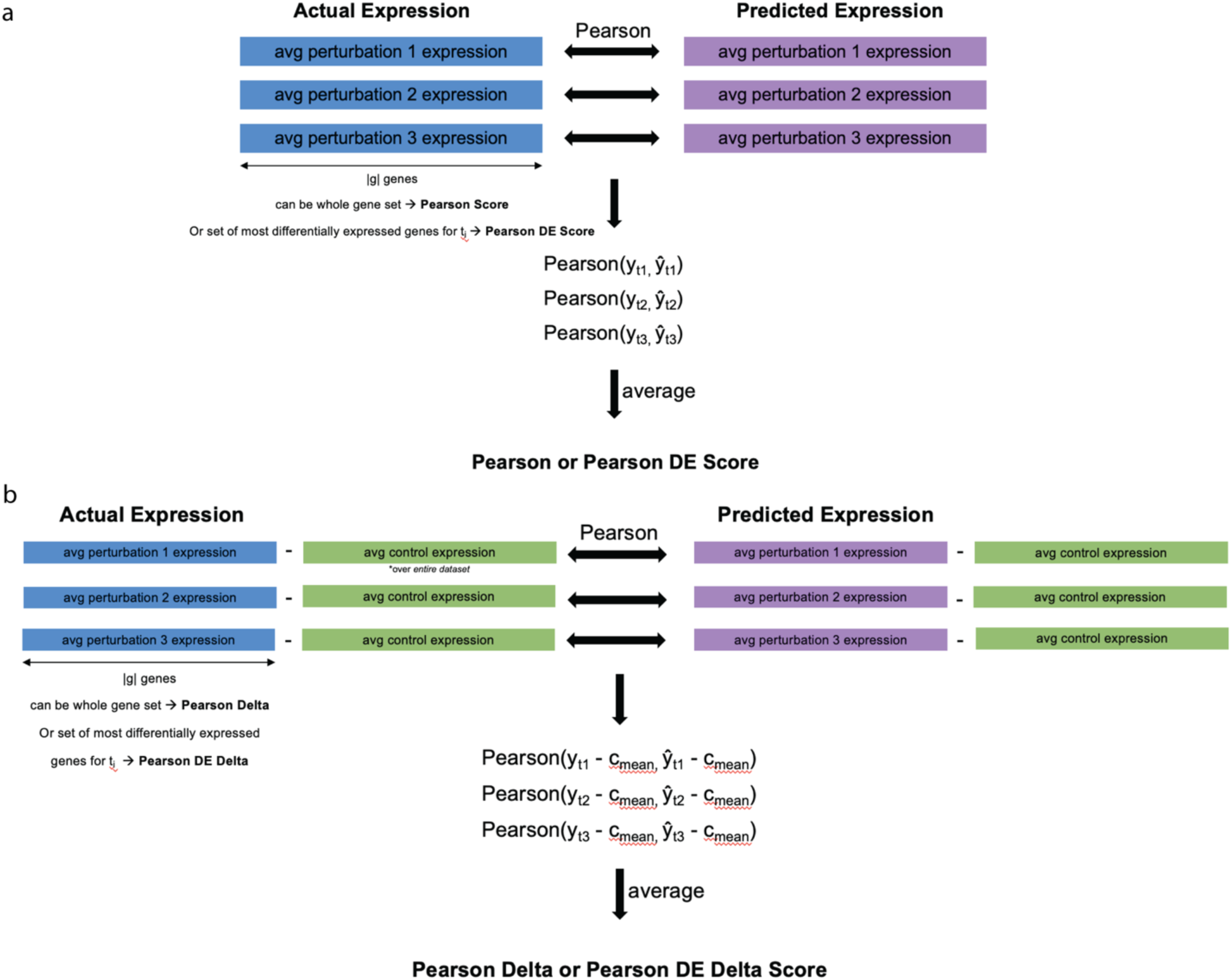
**Pictorial representation of Pearson metrics for measuring post-perturbation prediction performance.** (a) Pearson and Pearson DE scores. y_ti_ = actual expression of cells with target=t_i_. (b) Person Delta and Pearson DE Delta scores.

**Supplemental Figure 2:**
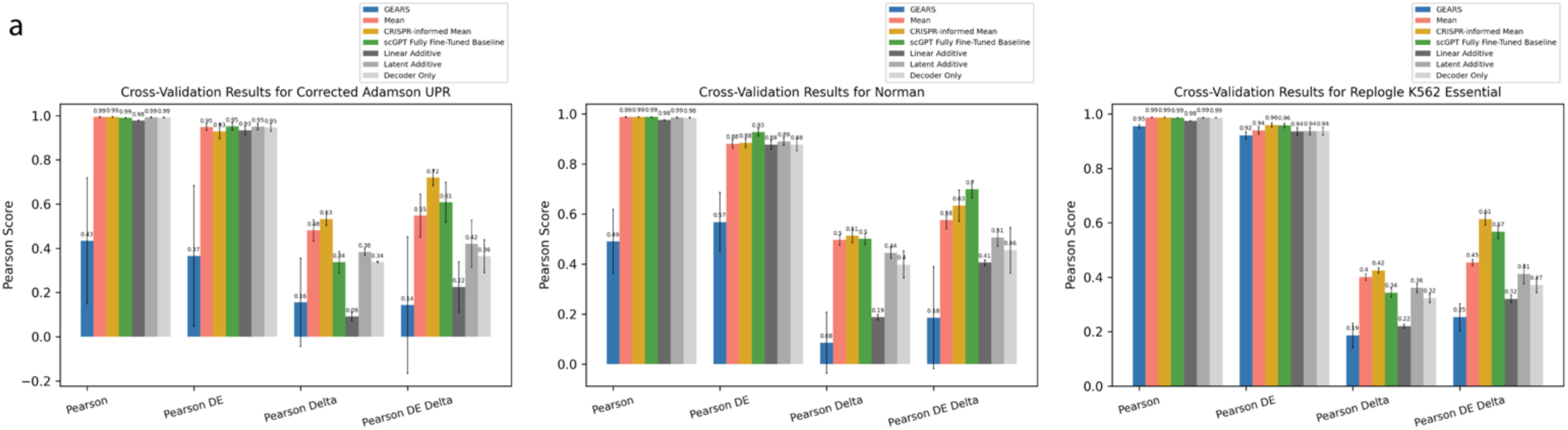
Four-fold cross-validation. We split the perturbations into four random folds, and assigned two of them for training, one for validation, and one for testing. Error bars show the standard deviation of the four folds.

**Supplemental Figure 3:**
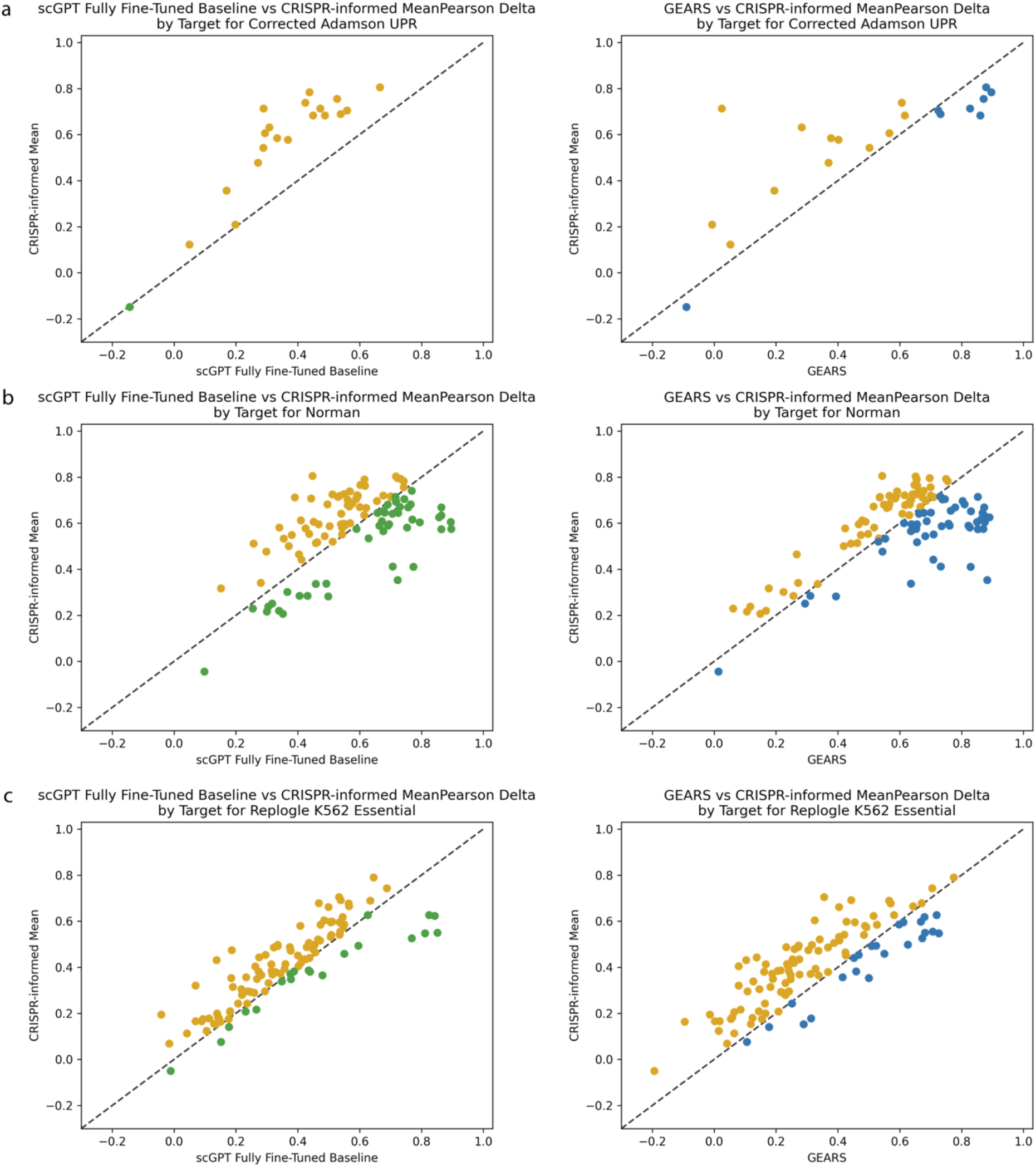
Performance by perturbation condition over the test set. X-axis DL model (either scGPT (left) or GEARS (right), Y-axis CRISPR-informed Mean Model. Each point is a perturbation condition. We used the best performing GEARS and scGPT models of the ten independent runs to derive predictions. The color indicates which model was the most performant for that condition: CRISPR-informed model: gold, scGPT: green, GEARS: blue. (a) Corrected Adamson UPR dataset. (b) Norman dataset. (c) Replogle K562 Essential dataset.

**Supplemental Figure 4:**
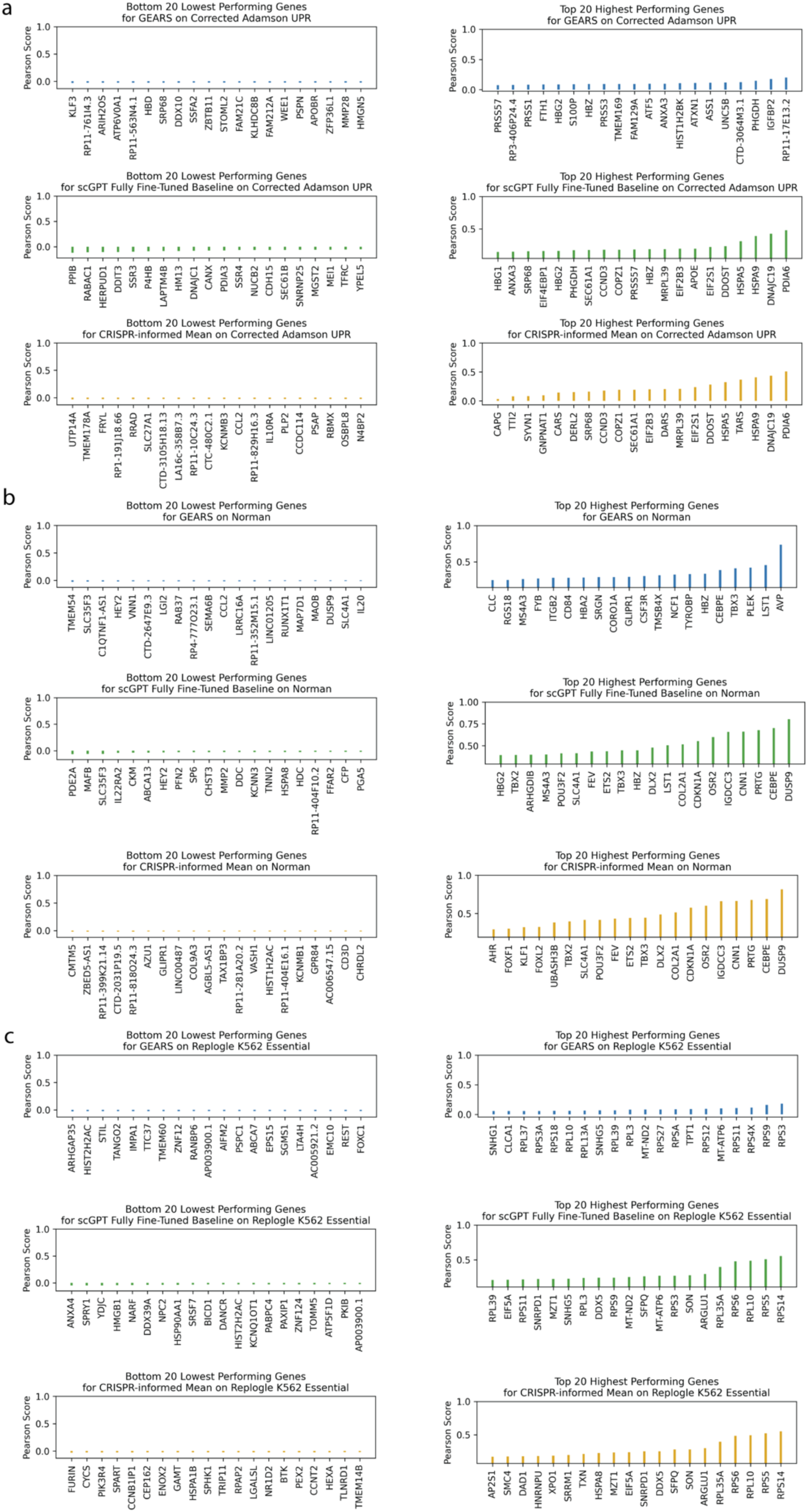
Performance by gene over test set. Left: Bottom 20 lowest performing genes for each method. Right: top 20 highest performing. We took the Pearson correlation between the predicted and the actual expression for all conditions in the test set. We used the best performing GEARS and scGPT models of the ten independent runs to derive predictions. For the CRISPR-informed mean model, the predicted expression for gene g is the same for every cell (by definition). This resulted in a prediction vector with zero standard deviation, and hence an invalid Pearson correlation. For this plot only, we added uniform random noise in the range 0 to 1×10^-7^ to each CRISPR-informed mean prediction. (a) Corrected Adamson UPR dataset. Gene overlap between top 20 of scGPT and CRISRP-informed mean: 12/20. (b) Norman dataset. Gene overlap between top 20 of scGPT and CRISRP-informed mean: 15/20. (c) Replogle K562 Essential dataset. Gene overlap between top 20 of scGPT and CRISRP-informed mean: 12/20.

**Supplemental Figure 5:**
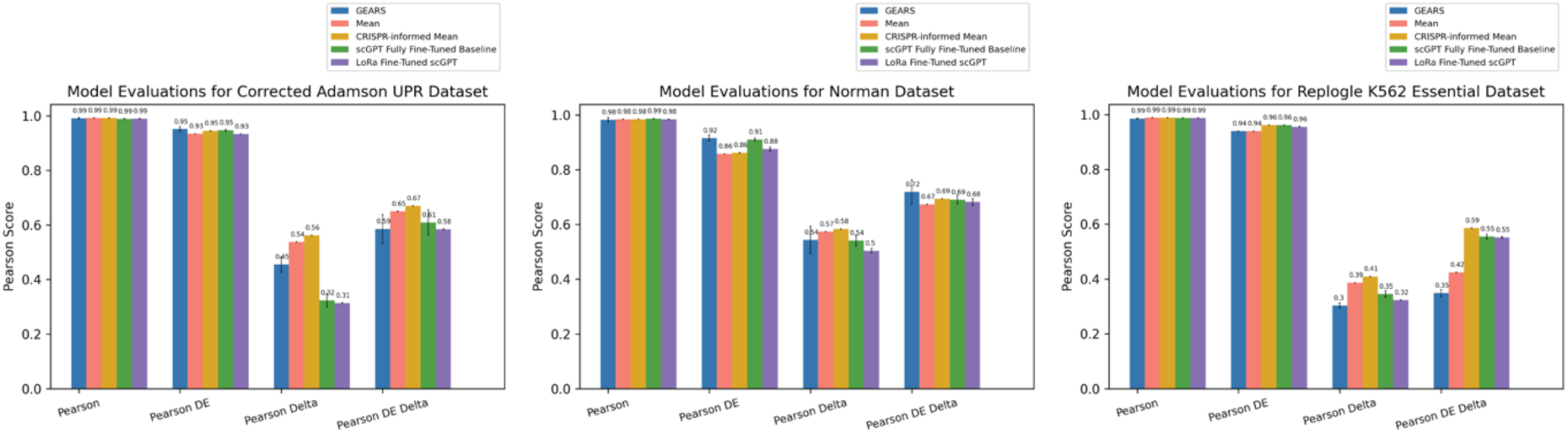
Performance bar charts on held-out test set comparing LoRa fine-tuned scGPT to other models. Error bars span plus and minus one standard deviation of ten independently trained models applied to the same test set.

**Supplemental Figure 6:**
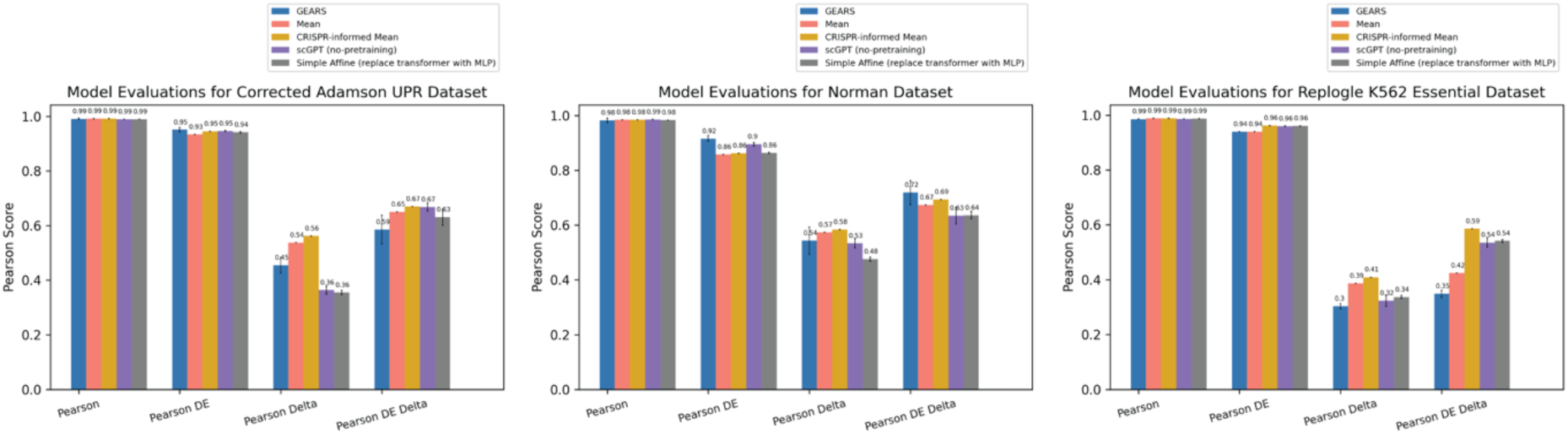
Simple Affine model with MLP replacement over test set. We replaced the transformer block of scGPT with a simple multilayer perceptron (MLP) that had the same number of layers as the original transformer block. The MLP used a ReLU non-linear transformation, with an equivalent embedding size as the transformer block (512). Error bars show the standard deviation across ten independent model runs.

**Supplemental Figure 7:**
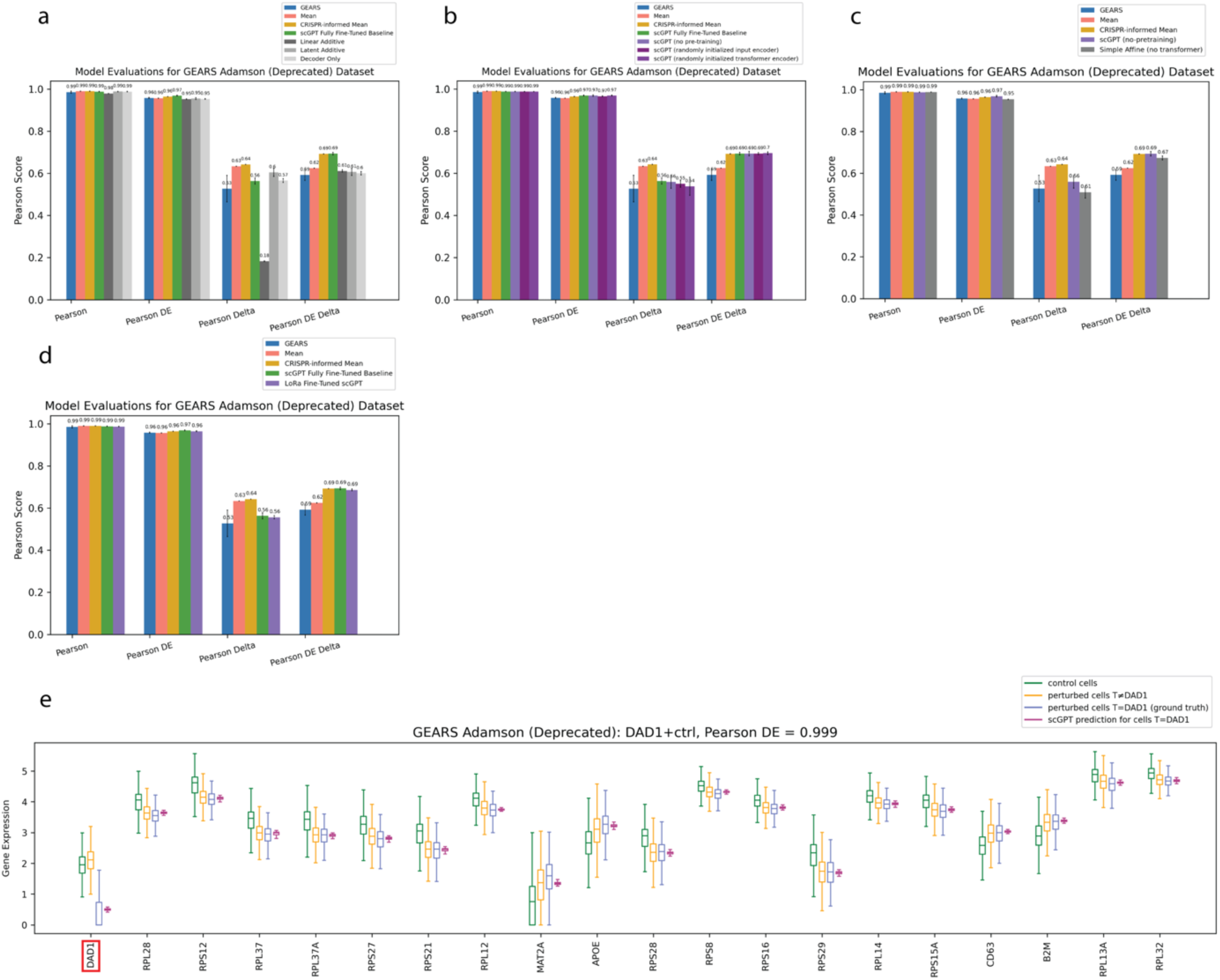
Results for the Deprecated GEARS Adamson dataset. (a) – (d) performance bar charts on held-out test set for different models. Error bars span plus and minus one standard deviation of ten independently trained models applied to the same test set. (a) Baseline performance across different models. (b) Various pre-training controls. (c) Simple affine performance. (d) LoRa fine-tuning. (e) Boxplots of top 20 DE genes for an example high-performing held-out perturbation condition (same perturbation as shown in the original scGPT paper^27^).

**Supplemental Figure 8:**
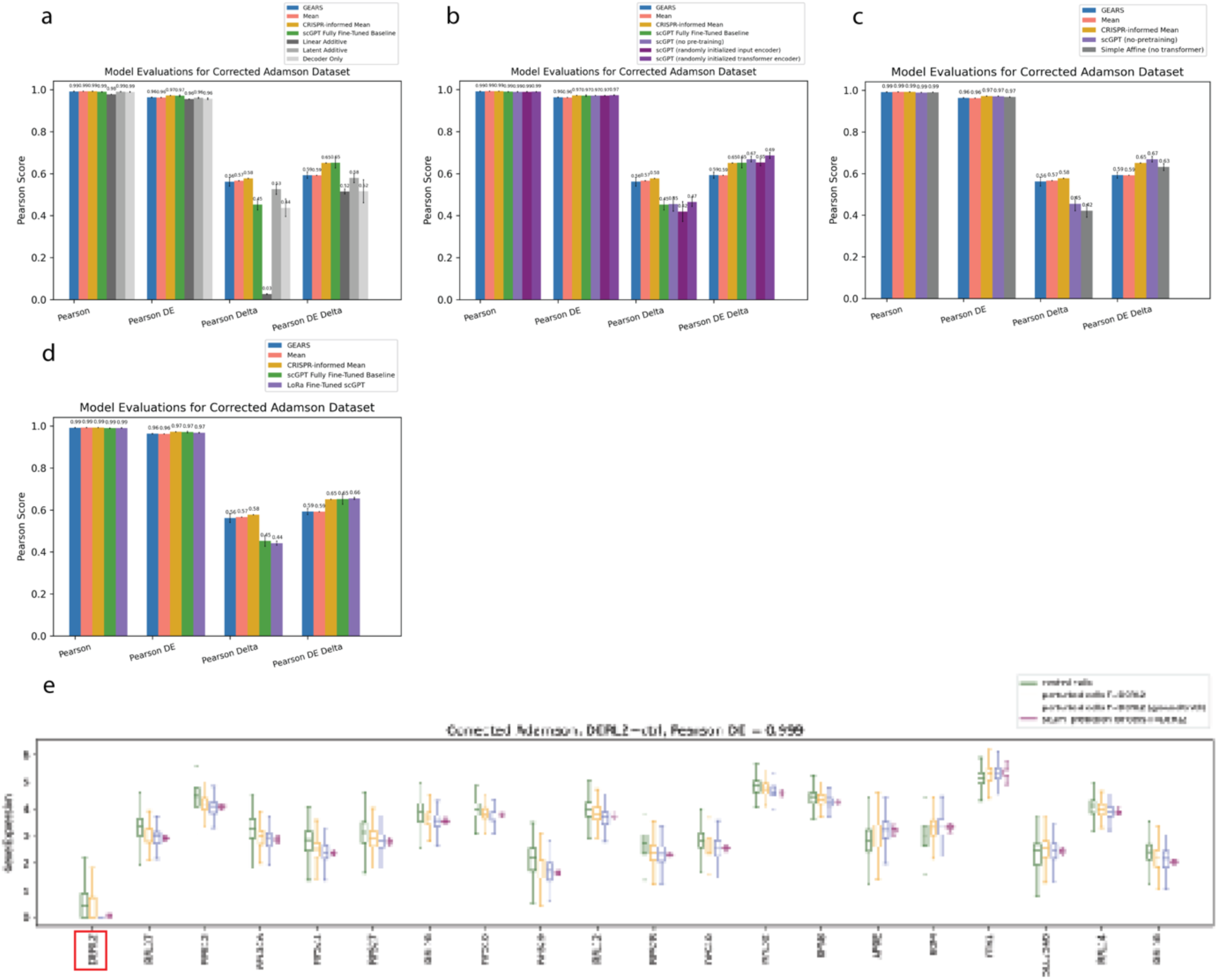
Results for the Corrected GEARS Adamson dataset. All models were trained on the corrected combination of the three sub-experiments. (a) – (d) performance bar charts on held-out test set for different models. Error bars span plus and minus one standard deviation of ten independently trained models applied to the same test set. (a) Baseline performance across different models. (b) Various pre-training controls. (c) Simple affine performance. (d) LoRa fine-tuning. (e) Boxplots of top 20 DE genes for an example high-performing held-out perturbation condition.

**Supplemental Figure 9:**
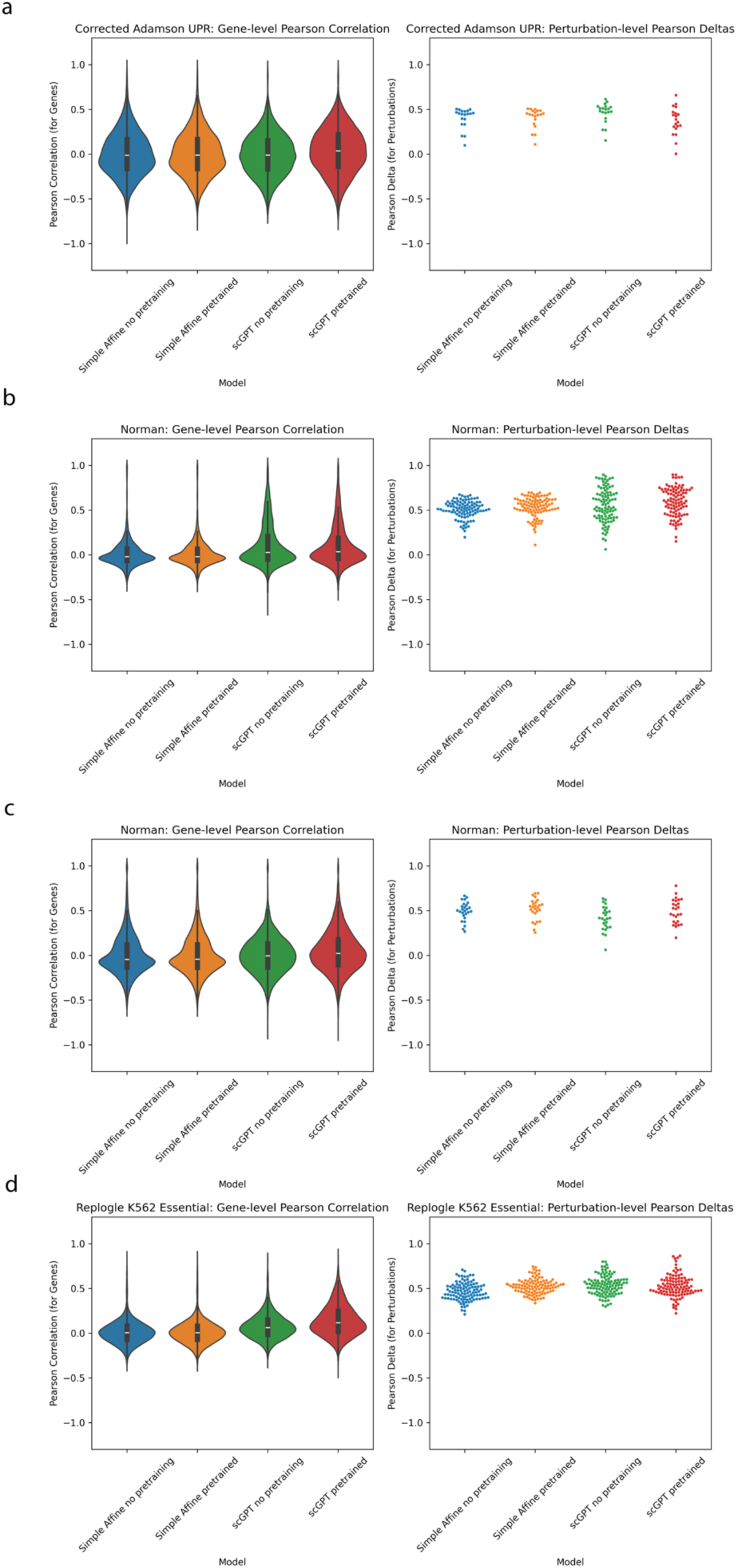
Comparing attention-based models to attention-free models with and without pre-training. Plots of gene-level Pearson correlations between actual condition means and predicted means (left) and perturbation-level PD (right). Violin plots were used when data points exceeded 500 points else swarm plots. Perturbation-level PD is calculated using the actual and predicted expression of all genes for a given condition. Gene-level Pearson correlations are calculated using the transpose of those matrices, with the correlation calculated between the actual and predicted expression of each gene across all perturbations. The target gene itself is excluded because it is a highly influential point and not directly relevant to the task of downstream biological predictions. X-axis indicates which model the predictions come from: (1) Simple Affine, (2) Simple Affine with pretrained input encoder weights, (3) scGPT architecture without pretraining and (4) fully fine-tuned scGPT. (a) Adamson UPR experiment data only (b) Norman single and double perturbation data (c) Norman just single perturbation data (the same gene can be present in both train and test for double perturbation data, but not single) (d) Replogle K562 essential dataset. There is an apparent improvement in predictions using scGPT compared to the other models. The best-predicted conditions are those for which ribosomal proteins (RP) are targeted (5 conditions in the test set), and the best-predicted genes are those that change in RP targeted conditions. This observed effect may be biologically relevant learning resulting from the combination of the scGPT architecture and the pre-training, but we cannot rule out the possibility that it could be the result of an unknown or unobserved technical artifact.

**Supplemental Figure 10:**
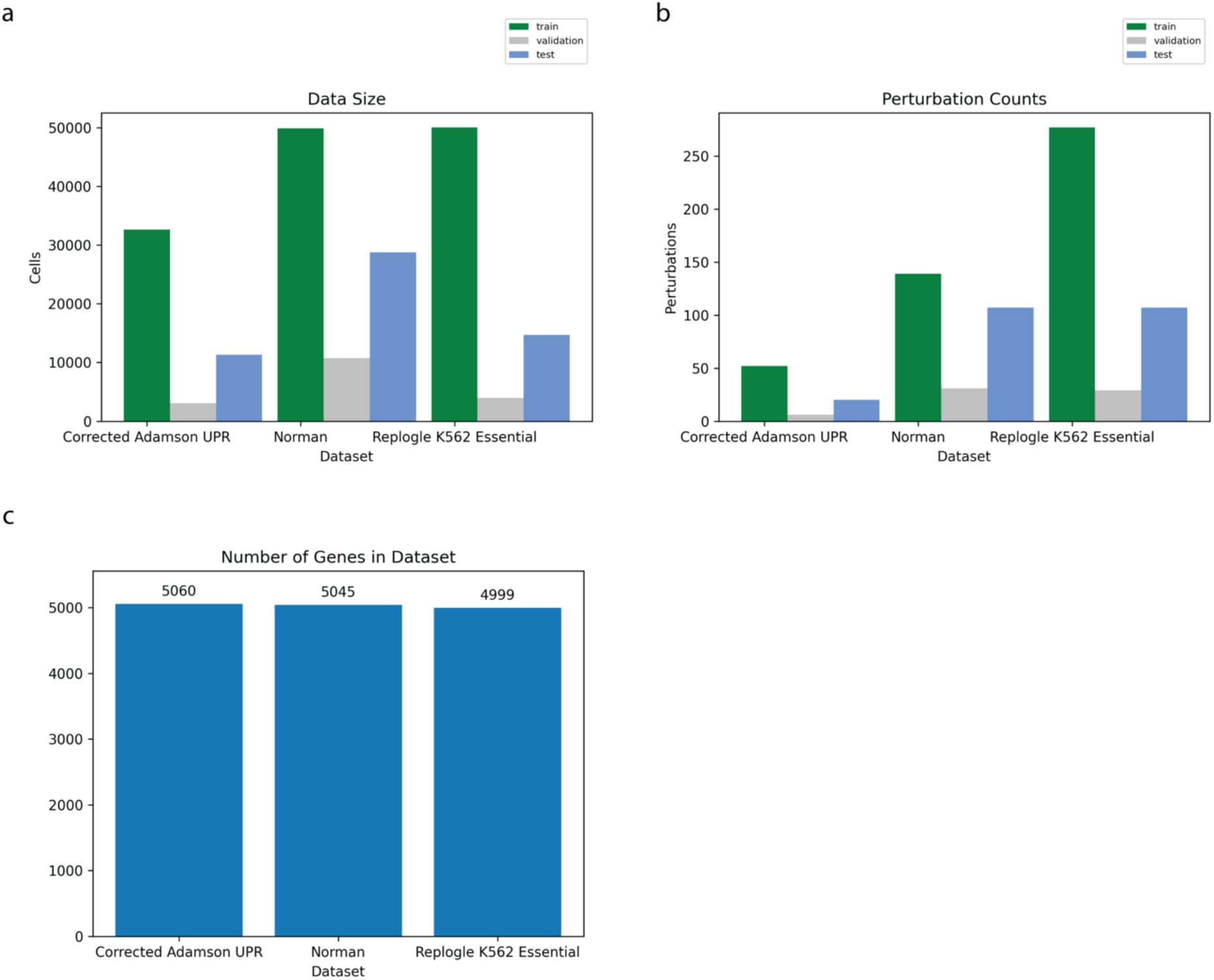
**Cell, perturbation, and gene counts.** (a) Cell counts for each dataset for each data split. (b) Unique perturbation counts. Combination perturbations in the Norman dataset are counted as unique. (c) Number of genes in each dataset.

